# Vpr R36W and R77Q Mutations Alter HIV-1 Replication and Cytotoxicity in T Lymphocytes

**DOI:** 10.1101/2020.05.26.118174

**Authors:** Antonio Solis-Leal, Dalton C. Karlinsey, J. Brandon Lopez, Vicente Planelles, Brian D. Poole, Bradford K. Berges

## Abstract

Chronic immune inflammation (CII) is a characteristic symptom of HIV-1 infection that contributes to acquired immunodeficiency syndrome (AIDS) progression in infected patients. Distinct AIDS development rates have shown that there are Rapid Progressor (RP) and Long-Term Non-Progressor (LTNP) patients, but the circumstances governing these differences in disease progression are poorly understood. Mutations in the Viral Protein R (Vpr) have been suggested to have a direct impact on disease progression. Studies have shown that Vpr interacts with both host and viral factors; these interactions affect cellular activities such as cell cycle progression and enhancement of apoptosis. The Vpr mutants R36W and R77Q have been associated with RP and LTNP phenotypes, respectively; however, these findings are still controversial. This study sheds light on the effects that Vpr mutations have in the context of HIV-1 infection of the HUT78 T cell line, using replication-competent CXCR4-tropic virus strains. Our results show a replication enhancement of the R36W mutant (increased viral load and percentage of p24+ cells) accompanied by increased cytotoxicity. Interestingly, the R77Q mutant showed a unique enhancement of apoptosis (measured by Annexin V and TUNEL staining) and G2 cell cycle arrest; these effects were not seen with WT or R36W viruses. Since necrosis is associated with the release of pro-inflammatory factors, the R36 mutation could lead to more robust CII and the RP phenotype. Conversely, the R77Q mutation leads to apoptosis, potentially avoiding CII and leading to a LTNP phenotype. Thus, Vpr mutations may impact HIV-1 related progression to AIDS.

**Importance:** The *vpr* gene is thought to be an important virulence factor in Human Immunodeficiency Virus type 1 (HIV-1). *vpr* polymorphisms have been associated with different rates of acquired immunodeficiency syndrome (AIDS) progression. However, there is controversy about the cytopathic and virulence phenotypes of Vpr mutants, with contradictory conclusions about the same mutants. Here, we examine the replication capacity, apoptosis induction, and G2 cell cycle arrest phenotypes of three *vpr* mutants compared to wild-type HIV-1. One mutant associated with rapid AIDS progression replicated more efficiently and killed cells more rapidly than wild-type HIV-1. Another mutant associated with slow AIDS progression triggered apoptosis more efficiently than wild-type HIV-1. These results shed additional light on the role of *vpr* polymorphisms in T cell killing by HIV-1 and may help to explain the role of Vpr in different rates of AIDS progression.

## Introduction

Vpr is found in many types of lentiviruses, including Human Immunodeficiency Virus type 1 (HIV-1), HIV-2 and Simian Immunodeficiency Virus. It has a critical role in pathogenesis and replication, and is classified as a viral regulatory protein (1, 2). The *vpr* gene has been shown to be highly conserved across many types of lentiviruses, suggesting the biological importance of Vpr in the viral life cycle (3, 4).

Although *vpr* is a late gene, the mature virion carries many copies of the protein inside the capsid, and thus it is active during early stages of infection (5, 6). Vpr carries out different functions during the viral infection cycle. It is part of the pre-integration complex, which facilitates the translocation of viral DNA to the nucleus of non-dividing cells (7). It also likely plays a crucial role in counteracting cellular antiviral factors; this activity is highly related to its morphology and interactions with other Vpr proteins or host factors (8). Furthermore, the presence of Vpr has been shown to enhance virus replication in T cells (9). This effect could stem from several of the interactions that Vpr has with the host cell, including Vpr-mediated G2 phase cell cycle arrest (10–13) or through transcriptional modulation (14–17). During G2 phase, the HIV Long Terminal Repeat promoter is more active (18), suggesting that viral-induced G2 arrest could be a tool to enhance viral replication.

One of the other functions most commonly associated with Vpr is the regulation of apoptosis in infected cells (19, 20). Early studies showed that Vpr can cause the permeabilization of the permeability pore complex (PTPC), resulting in the release of apoptotic factors from the mitochondrial membrane (20–22). However, it was later shown that Vpr-induced depolarization of the mitochondrial membrane was not a result of Vpr binding to PTPC, but rather due to activation of the Bax protein, which is a result of stress signals sent by the Ataxia Telangiectasia and Rad3-related (ATR) DNA damage response pathway (23). The ATR pathway responds to DNA damage, and when activated it initiates a signaling cascade that stalls the cell at a cycle checkpoint (24). Activated ATR has also been shown to phosphorylate Breast Cancer type 1 susceptibility protein (BRCA1), which in turn leads to the upregulation of Growth Arrest and DNA Damage inducible alpha protein (GADD45α), which eventually leads to Bax activation and the release of apoptotic factors from mitochondria. Knocking down ATR expression with RNAi abrogated both the G2 arrest and apoptosis associated with Vpr expression, showing that the processes happen concomitantly (23–25). It has been hypothesized that ATR is activated in response to Vpr-induced herniations in the nuclear membrane (25, 26). Despite both processes being regulated by ATR activation, Vpr induction of G2 arrest and apoptosis are independent processes (18, 27–30). It has been observed that HIV-infected cells are more commonly apoptotic during G1 or S phase, while cell cycle arrest occurs in M or G2 phase (22, 27, 30, 31). Both virion-associated and *de novo*-synthesized Vpr are able to induce apoptosis and cell cycle arrest through identical pathways (27, 32). Additionally, secreted Vpr has been suggested to cause apoptosis by the permeabilization of cellular membranes to calcium and magnesium (33–35). Vpr- and reverse transcription-induced apoptosis have been shown to be the main causes of cell death in peripheral blood resting CD4+ T cells, potentially contributing to bystander cell death associated with HIV infections (36).

Acquired immunodeficiency syndrome (AIDS) is a consequence of the tropism of HIV, which targets CD4+ T cells. These cells are essential for coordinating adaptive immune responses to pathogens, but HIV infection can cause a decrease in their concentration in blood to less than 200 CD4+ T cells/μL of blood, compared to about 1,000 CD4+ T cells in healthy individuals (37). Depletion of CD4+ T cells is commonly observed in HIV patients. Patients that maintain normal levels of CD4+ T cells for many years are classified as long term non-progressors (LTNP) while those that experience a rapid decrease in CD4+ T cells are classified as rapid progressors (RP); RP patients usually develop AIDS 5 to 8 years after infection (38). Mutations in Vpr have been proposed as a causative agent of the phenotypic differences observed in infected patients (39, 40), but there is much that is not understood due to the complexity of both viral and host genetics. Various mechanisms have been proposed to explain how Vpr could enhance disease progression: A) Promoting early T cell activation, by enhancing Nuclear Factor of Activated T cells (NFAT) to prime non-activated T cells for productive infection (41); B) Inhibiting T cell proliferation and enhancing cell death (3); or C) Inducing the infection and activation of virus production from latency in macrophages, creating drug-resistant HIV reservoirs (42–44).

The Vpr R77Q mutant has been suggested to produce a LTNP phenotype in HIV-infected patients. This mutation has been found in several LTNP patients, and has also been reported to produce low virus replication (9, 39, 45). However, other studies have shown that this mutation does not confer a clinical advantage to subjects receiving antiretroviral therapies (46, 47), increasing the controversy around the cytopathogenicity of this mutation. On the other hand, the Vpr R36W mutation has been associated with rapid progression, but was shown to have similar replication and CD4+ T cell depletion rates to wild-type strains (WT) (9). Vpr R36W mutants also may have an enhanced capacity to oligomerize that could impair its ability to induce G2 cell cycle arrest and enhance disease progression (9).

In this study, we analyze the replication rate, G2 arrest, and apoptotic phenotypes of HUT78 cells infected with either HIV-1 expressing either wild-type Vpr, R36W, R77Q, or Vpr null mutants to better understand the role that Vpr plays in viral replication and T cell killing pathways as they relate to disease progression.

## Results

### Construction of *vpr* mutants

All virus strains used in this study were based on the wild-type (WT) pNL4-3 molecular clone, which uses CXRC4 as a co-receptor. The R36W and R77Q constructs were produced by replacing the WT *vpr* gene with mutated *vpr* sequences from expression plasmids obtained from Dr. Velpandi Ayyavoo (9). We created the null mutant to study the effects of HIV infection with a defective *vpr* gene but without any mutations in neighboring genes/reading frames. This was accomplished using site-directed mutagenesis (SDM) to mutate the *vpr* start codon (ATG to GTG, or M1V), resulting in a silent mutation in nucleotide 173 of the overlapping *vif* gene (AGA to AGG; both encode arginine).

Figure 1A shows the differences in nucleotide sequences between each strain and the corresponding change in amino acid sequence. Sanger DNA sequencing was used to confirm the desired mutations and to show that no additional mutations were introduced during the creation of constructs (data not shown).

**Figure 1:**
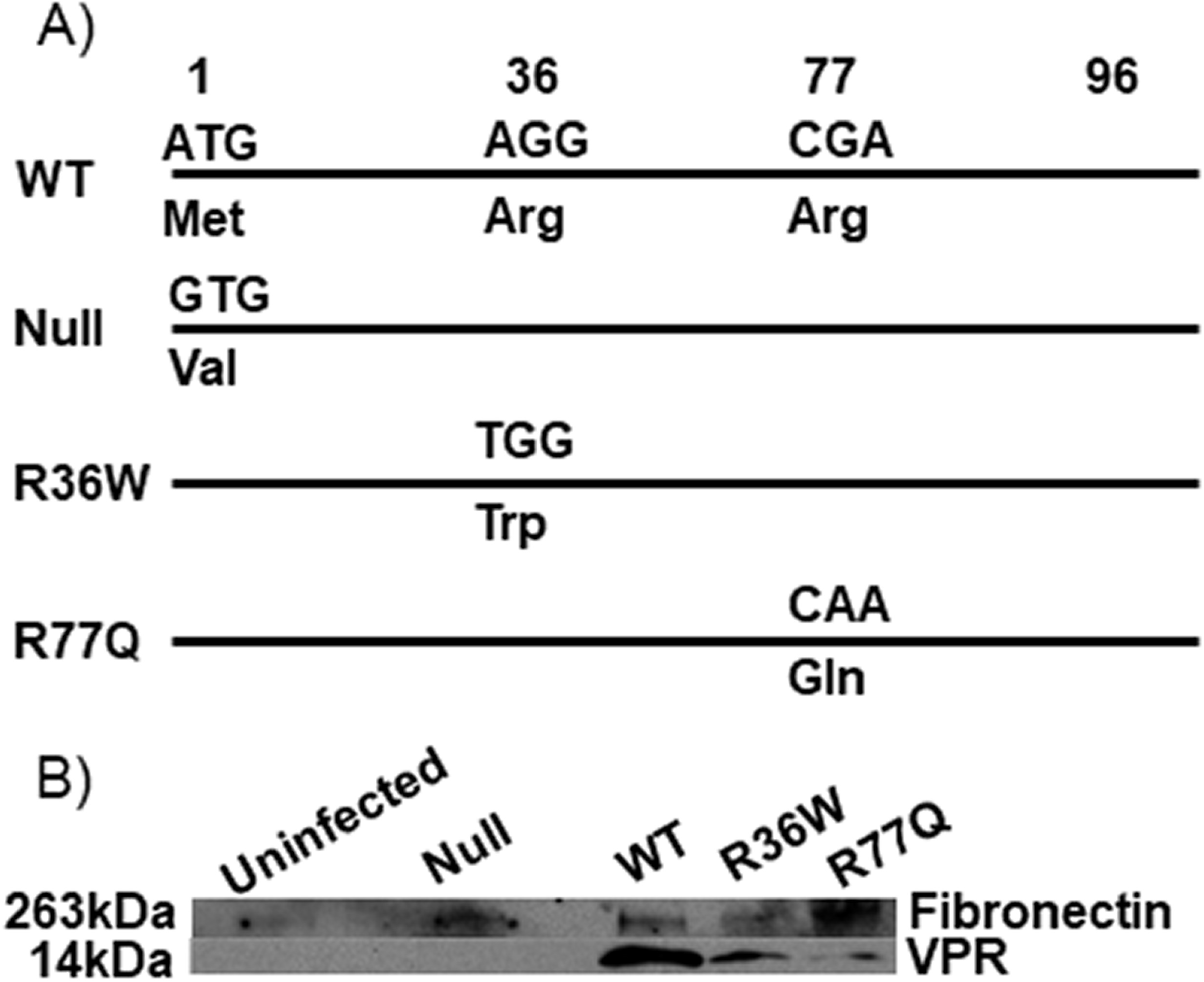
Differences in nucleotide and amino acid sequences between *vpr* mutants and confirmation of proper expression profiles by immunoblot. **(A)** Sequence comparison of the mutations produced by site-directed mutagenesis in NL4-3 HIV *vpr*. **(B)** Immunoblotting confirmed Vpr expression in HUT78 cells infected with WT, R36W, or R77Q while no Vpr expression was detected in the null mutant or in uninfected cells. Fibronectin was used as a loading control.

To confirm the expression of Vpr in WT, R36W and R77Q mutants, and the lack of expression in the null mutant, we detected Vpr expression via immunoblotting. Infected HUT78 cell lysates collected at 7dpi at MOI 0.01 were used. We found that WT virus as well as the R36W and R77Q mutants produced detectable Vpr, while no Vpr was detected in the null mutant or in uninfected cells (Fig. 1B).

### The R36W mutant has an enhanced replication capacity relative to WT virus

To examine the replicative capacity of our mutants in a T cell line, HUT78 cells infected at 0.01 MOI were analyzed for viral replication via Q-RT-PCR over a 7-day period post-infection (Figure 2A). After 3 days, the only statistically significant difference in genome copy number that we observed was between the R77Q and WT populations. The R77Q infected population had a viral load of 1.0×10^8^ RNA copies/mL, which was almost tenfold higher than the load of the WT virus (1.7×10^7^ copies/mL), but at 5 and 7 days post-infection (dpi), there was no difference between the two populations. The R36W-infected population had a ten-fold higher load (2.9×10^9^ copies/mL) compared to all other strains at 5 dpi; while still statistically significant 7 dpi, the difference between R36W and the other strains was considerably smaller (Figure 2A). There was no significant difference in viral load between the WT, R77Q, and null viruses at either 5 or 7 dpi (Data Set S1).

**Figure 2:**
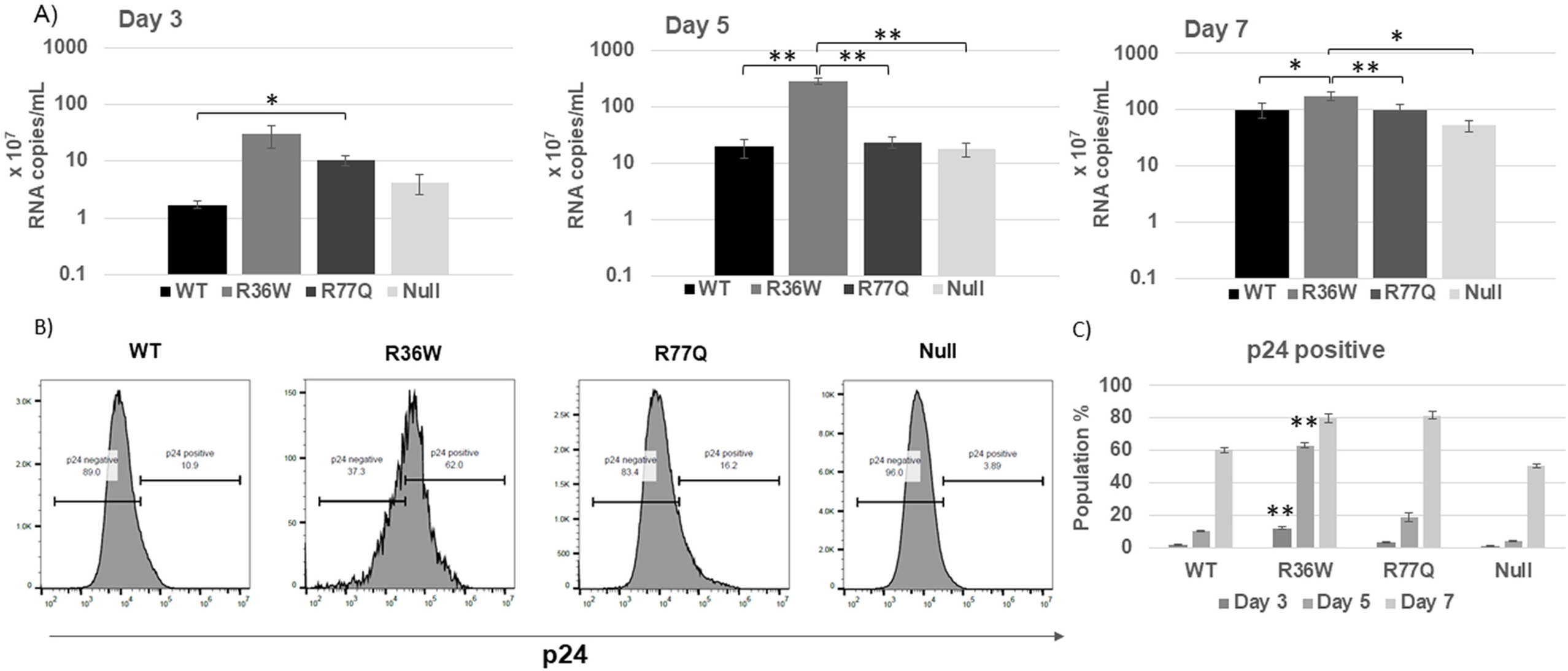
R36W mutant shows an enhanced viral replication and a rapid formation of a p24 positive population. R36W, R77Q or null HIV Vpr mutants, or WT NL4-3 were used to infect HUT78 cells at MOI 0.01. At 3, 5 and 7 dpi supernatants were analyzed for viral load and cells were analyzed for p24 expression. **A)** Quantification of viral replication was determined through Q-RT-PCR. **B)** Assessment of p24+ populations via flow cytometry. Histograms show representative results from 7 dpi. In **B**, asterisks only show statistical differences between R36W compared to all other samples. The data are representative of 3 independent experiments and error bars indicate Standard Error (SE). * p-value ≤ 0.05, ** p-value ≤ 0.01. See supplementary material S1 and S2 for complete statistical analysis.

Each mutant’s capacity to express viral genes was studied by measuring the p24 expression of individual cells through flow cytometry (Figure 2B/2C). Cells infected with the R36W mutant showed a significantly increased p24+ population at all three time points compared to the WT and null viruses (Data set S2). R36W also had much higher p24 expression than R77Q at 3 and 5 dpi, but on day 7, both populations were about 80% p24+. The mean fluorescent intensity (MFI) of p24+ histograms was also examined and showed that p24 production in individual cells was also higher following infection with the R36W mutant (Data Set S3).

### R77Q mutation leads to an increase in the percentage of apoptotic cells

To test whether the Vpr mutants induced differing levels of apoptosis, we infected HUT78 cells at 0.01 MOI and measured Annexin V and Fixable Viability Dye (FVD) binding, via flow cytometry._Annexin V binds to phosphatidylserine, which flips from the inner layer of the plasma membrane to the outer layer during the early stages of apoptosis (48). Cells were also stained with FVD, which indiscriminately binds proteins. It enters dead cells through holes in the plasma membrane, causing them to fluoresce with increased intensity compared to live cells. Since cells with a permeabilized membrane will allow Annexin V to bind phosphatidylserine inside the plasma membrane, we considered cells positive for both FVD and Annexin V to be necrotic while cells positive for Annexin V only were considered to be apoptotic.

The results of this assay (Figure 3) showed a clear difference in apoptosis induction by the R77Q mutant. At all three time points, the R77Q-infected population had significantly higher levels of apoptosis; at 7 dpi, the R77Q population had a mean level of 47.4% Annexin V single-positive cells, compared to 2.64% in WT, 5.54% in R36W, 7.26% in null, and 3.29% in the uninfected population (Figure 3A/3B). The only significant difference in the number of apoptotic cells in the WT and R36W populations was 7 dpi; on days 3 and 5, the two populations had similar numbers of apoptotic cells (Data Set S4B). The null population also had a significant increase in the number of apoptotic cells (7.26%) at 7 dpi compared to the WT population. The null population had similar numbers of apoptotic cells to R36W at both 3 and 7 dpi, but interestingly had a significantly (p=0.021) lower number of apoptotic cells when observed 5 dpi (Data Set S4B). The number of apoptotic cells in the WT population was not significantly different from the uninfected population at all three time points.

**Figure 3:**
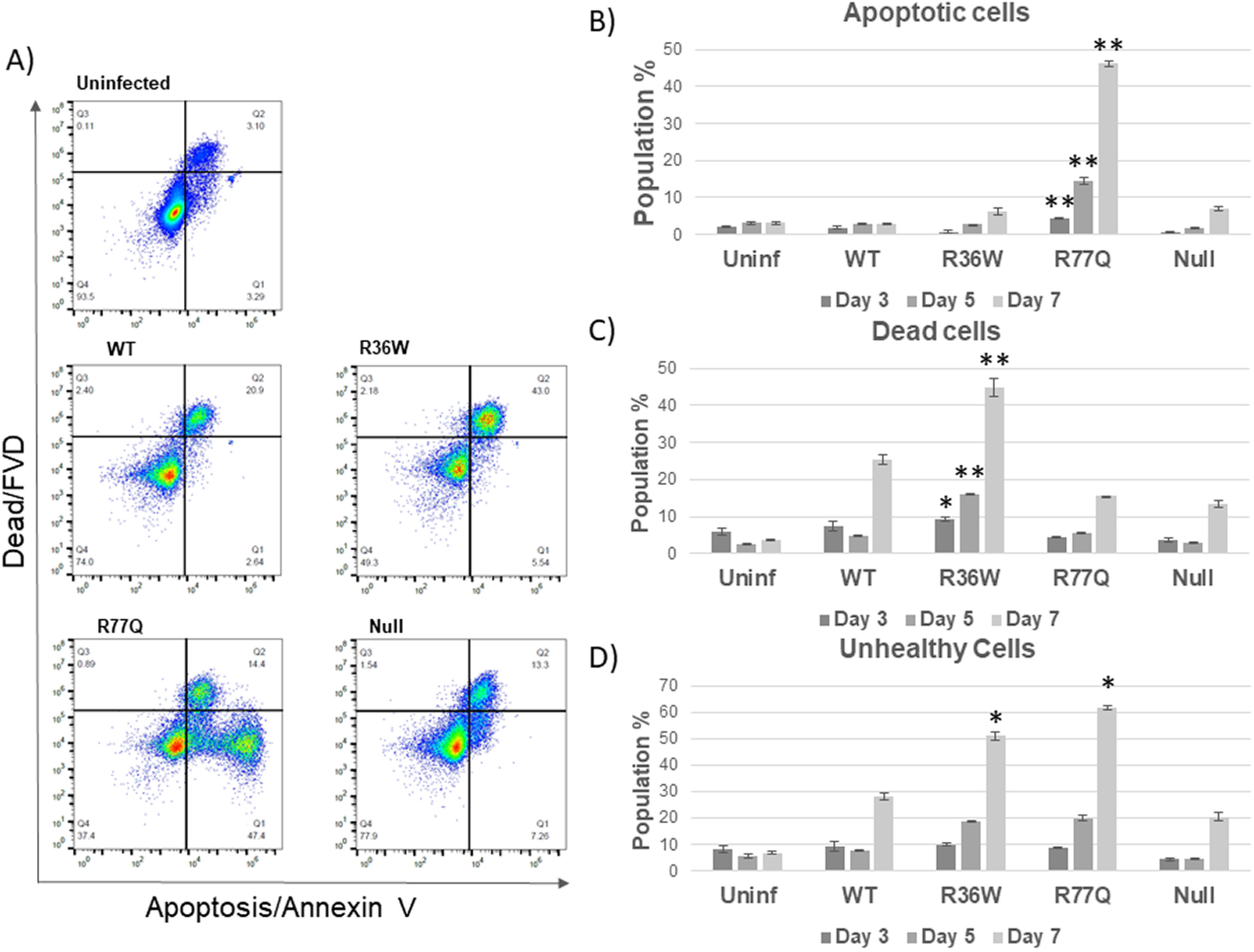
R36W and R77Q Vpr mutants trigger different death pathways. R36W, R77Q or null HIV Vpr mutants, or WT NL4-3 were used to infect HUT78 cells at MOI 0.01. Cell samples were analyzed at 3, 5 and 7 dpi. Apoptosis was detected by Annexin V staining and fixable viability dye (FVD) was used to detect dead cells. **A)** Representative dot plots from samples collected on day 7 pi. **B)** Apoptotic cells (positive for Annexin V only). **C)** Dead cells (all cells positive for FVD). **D)** Unhealthy cells (all cells staining positive for FVD and/or Annexin V). Asterisks show statistical differences between R36W (panels C and D) or R77Q (panels B and D) and all the other samples. Data are representative of 3 independent experiments and error bars indicate SE. * p-value ≤ 0.05, ** p-value ≤ 0.01. See supplementary material S4 for complete statistical analysis.

We also observed clear differences in the number of dead cells in each population. At 7 dpi, all four infected populations had significantly higher numbers of dead cells than the uninfected control (Figure 3C, Data Set S4A). The percentage of dead cells in the R36W- and WT-infected populations was significantly higher than in the R77Q and null populations. Both 5 and 7 dpi, there was a significant difference between the R36W and WT populations. At both time points, the number of dead cells in the R36W population was twice as high as in the WT population (Figure 3A/3C). The percentage of dead cells in the null-infected population was significantly smaller than the other infected populations; it had relatively similar levels of apoptosis to the WT but fewer overall dead cells.

To get a better sense of the overall health of each sample population, we combined all the FVD and Annexin V positive populations into a single “unhealthy” cell population (Figure 3D). At 3 dpi the R36W population had a small 1.16% increase in the number of unhealthy cells compared to R77Q (p=0.035). Although this difference is significant, R36W and R77Q values were not statistically different to the WT unhealthy population on the same day (Data Set S4C). At 5 and 7 dpi however, the R36W and R77Q populations had around twice the number of unhealthy cells as the WT population. At 7 dpi, over 60% of the R77Q population was unhealthy compared to around 50% in the R36W population and almost 30% in the WT population (Figure 3D). The lack of an FVD (-), Annexin V (+) population in our results (Figure 3A) suggests that the WT and R36W mutants either induce extremely rapid apoptosis, or that they preferentially activate a necrotic pathway to kill cells. These data suggest that the R36W and R77Q mutants are significantly more cytopathic than the WT virus, but that they cause cell death via different pathways.

### Vpr mutants induce different types of cell death

To confirm that the double-positive dead cells observed in the Annexin V staining (Figure 3A) had not simply undergone apoptosis more rapidly, infected cells were stained for DNA fragmentation, a classical sign of late-stage apoptosis (48). TUNEL analysis by flow cytometry showed minimal signs of DNA fragmentation in the WT, R36W, and null mutant infected populations at both 5 and 7 dpi, while the R77Q-infected population had a significantly increased percentage of apoptotic cells (Figure 4A/4B, Data Set S5). At 7 dpi, the R77Q-infected population was 8.34% positive for TUNEL, while all the other populations had minimal TUNEL+ cells. Figure 4C shows a comparison of our apoptosis results as detected by either Annexin V or TUNEL staining. The R77Q mutant showed a significantly higher proportion of cells with apoptotic markers on days 5 and 7 compared to all other samples, as detected by each assay. The percentage of R77Q-infected cells positive for Annexin V was higher than the percentage of cells positive for TUNEL at days 5 and 7. This may be due to the fact that phosphatidylserine exposure occurs earlier during the apoptotic process than DNA fragmentation (48), allowing a greater portion of the apoptotic population to be identified. Together these data show that the R77Q mutant primarily kills cells through the induction of apoptosis, while suggesting that the WT, R36W, and null viruses cause death through different, non-apoptotic pathways.

**Figure 4:**
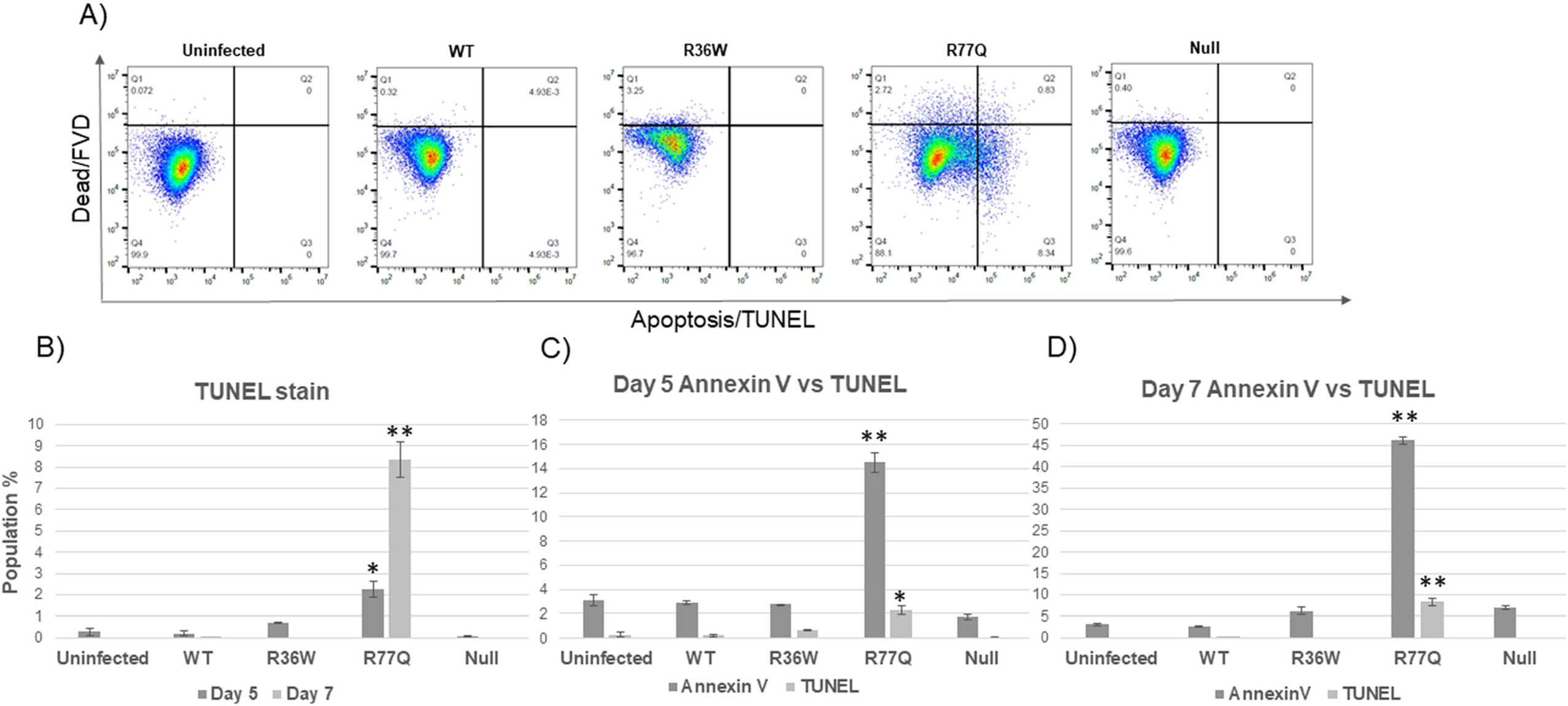
R77Q enhanced apoptosis confirmed by TUNEL stain. R36W, R77Q or null HIV Vpr mutants, or WT NL4-3 were used to infect HUT78 cells at MOI 0.01. Cell samples were analyzed at 5 and 7 dpi. **A)** Representative dot plots from day 7 pi. **B)** Bar graph representing the TUNEL single-positive cells of each sample. **C)** Bar graphs comparing apoptosis results obtained through TUNEL and Annexin V experiments on day 5 and 7. Asterisks show statistical differences between R77Q and all the other samples. Data are representative of 3 independent experiments and error bars indicate SE. * p-value ≤ 0.05, ** p-value ≤ 0.01. See supplementary material S5 for complete statistical analysis.

### G2 cell cycle arrest is most prominent in cells infected with the R77Q mutant

To determine if these mutants differed in their capacity to arrest cells in G2 phase, we used flow cytometry to observe the cell cycle progression of infected cells. HUT78 cells were infected with 0.1 MOI, then at 7 dpi cells were stained with propidium iodide and with antibodies to the p24 antigen and analyzed through flow cytometry. p24+ populations were gated to study cell cycle arrest specifically in infected cells. Only results from 7 dpi are reported due to a need for large numbers of p24+ cells for the analysis. Figure 5 shows a significant mean increase in detection of cells in G2 phase in most infected samples (WT=26.0%, p=0.01; R77Q=40.4%, p=0.01; null=26.0%, p=0.03) compared to the uninfected control (19.2%) (Data Set S6). R77Q showed the highest percentage of cells arrested in G2, which was significantly higher than the other infected samples (vs WT p=0.02, vs R36W p=0.04, vs null p=0.01). R36W G2 arrest was lower than the WT and not significantly different than uninfected. These results support studies that show that the R77Q mutant enhances cell cycle arrest and that R36W has a decreased capacity to cause cell cycle arrest (9). An analysis was also performed of p24-cells, but there was no significant difference of either WT Vpr or any mutants compared to uninfected controls (data not shown).

**Figure 5:**
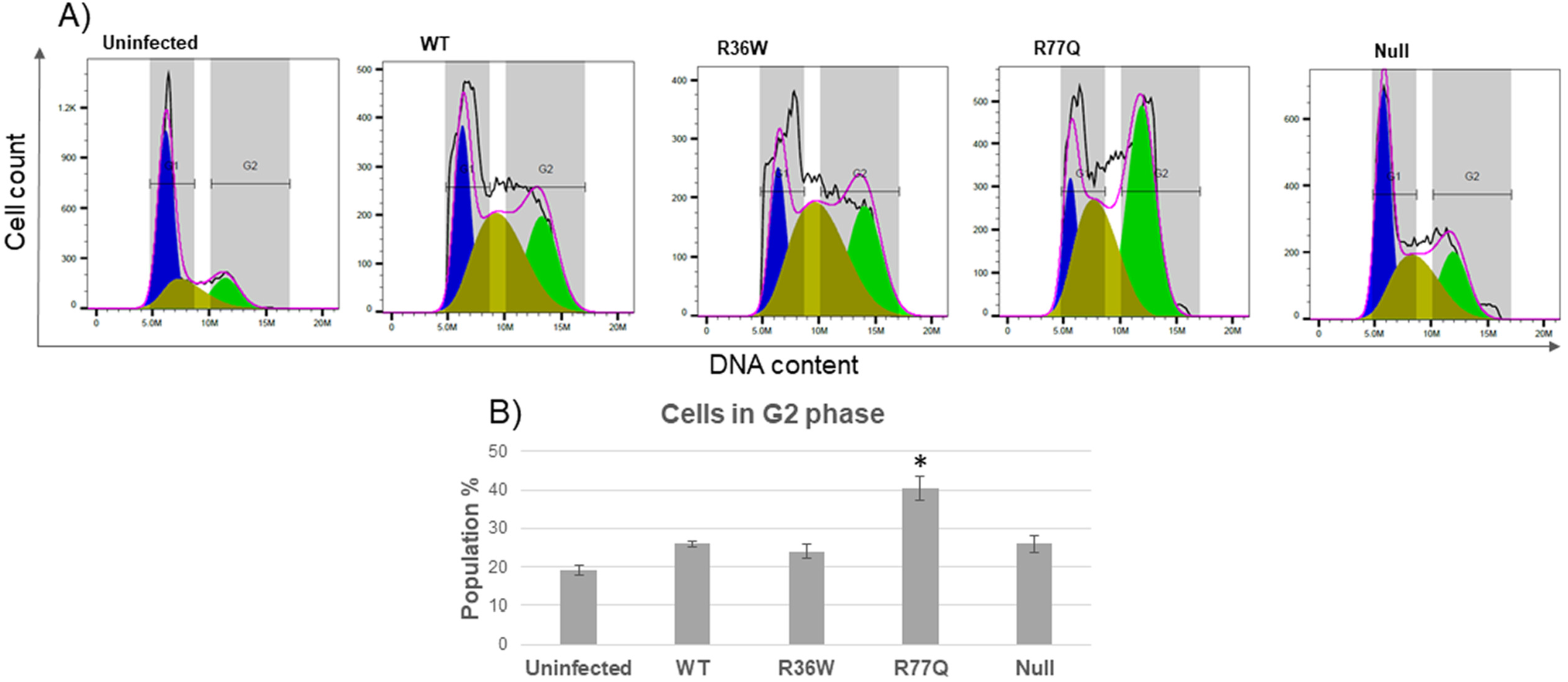
R77Q strongly enhances cell cycle arrest. R36W, R77Q or null HIV Vpr mutants, or WT NL4-3 were used to infect HUT78 cells at MOI 0.1. Cell cultures were gated on p24+ cells for cell cycle analysis. **A)** Representative histograms from day 7 pi. **B**) Percentage of p24+ cells in G2 phase. The asterisk shows a statistical difference between R77Q and all other samples. Data are representative of 3 independent experiments and error bars indicate SE. * p-value ≤ 0.05. See supplementary material S6 for complete statistical analysis.

## Discussion

In our efforts to study the phenotypes of the R36W and R77Q Vpr mutants to gain a better understanding of the role that HIV Vpr plays in disease progression; we have made four main conclusions: 1) The R36W Vpr mutation enhances the replicative capacity of HIV in HUT78 cells. 2) The R77Q Vpr mutation enhances the pro-apoptotic activity of HIV. 3) The WT, R36W and null viruses kill host cells via a necrotic, non-apoptotic pathway. 4) The R77Q Vpr mutation enhances G2 cell cycle arrest.

In our analysis of replication capacity by Q-RT-PCR of viral load and measurement of p24 expression via flow cytometry, we found that R36W-infected cells produced significantly more viral copies and p24+ cells compared to other strains. Our findings show that both the R77Q and the null mutants behaved very similarly to the WT virus in terms of viral replication rates, suggesting that these mutations do not impact viral replication efficiency in an *in vitro* setting. The mechanism by which the R36W mutant replicates more effectively in cell lines is still to be determined. It appears to be a gain of function mutation since this virus replicated more effectively than either WT or null virus, which showed similar replication efficiencies.

There was not a significant increase in the number of viral genome copies or p24+ cells in the R36W-infected population from days 5 to 7. This may be because the R36W-infected population had reached a plateau in terms of both HIV production and cell infection due to the limited number of infectable cells in the culture. As a result, the differences in the replication kinetics of each mutant were most visible at 5 dpi, and by 7 dpi the populations infected by the other mutants had a similar viral load. Overall, these results show that R36W has a significantly higher replication capacity compared to the WT virus. A previous study by Hadi et al. reported that WT virus showed the highest replication, followed by R36W (among other mutants studied) and that the R77Q mutant replicated at levels seven-fold lower than the WT virus (9). However, Hadi et al. constructed their mutants using a modified strain of HIV-1 NL4-3 engineered to express the EGFP reporter that places Nef expression under the control of an IRES element, which could result in attenuation (49). Therefore, cells infected with this virus may show different behavior than that observed with unmanipulated HIV-1 NL4-3. Much of the controversy regarding Vpr function is likely a result of the many different experimental systems used to study its effects including primary cells, cells in a three-dimensional lymphoid tissue, and cell lines. The Hadi et al study was conducted in primary cells, which could potentially vary in activation status, influencing the replication kinetics of the different viruses. Our use of a clonal population eliminates this variable.

The R77Q mutant showed significantly higher levels of G2 cell cycle arrest compared to WT virus and the other two mutants, in agreement with what has been previously reported by Hadi et al (9). Since the LTR promoter is known to be most active during G2 phase (50), but the R36W mutant showed both the lowest levels of G2 arrest and the highest replication rate, these results appear to be conflicting. Our p24 MFI results show a peak in p24 expression levels at day 7 for R77Q, which may support enhancement of viral gene expression due to LTR activation. Our viral load and p24 expression results show that the R36W virus still replicates significantly better in HUT78 cells, suggesting that the R36W mutation provides a gain of function that has a stronger effect on viral replication than the G2 arrest observed in R77Q infections.

A potential difference in these mutants that could affect disease progression is their distinct mechanisms of causing cell death. Contrary to the results of previous studies (23, 39), we observed much higher rates of apoptosis in the R77Q-infected population as compared to either the WT or the other mutant populations. The difference between our observed R77Q apoptosis phenotype and those previously reported is also likely a result of different experimental systems. The previous data that associated R77Q Vpr with a normal or decreased capacity for apoptosis was observed in cells transduced with R77Q Vpr-expressing plasmids (23) or in Jurkat cells infected with VSV-G pseudotyped virus (39). Our experimental system most closely resembles the natural conditions of infection since we have studied Vpr function in the context of replication-competent HIV, including the apoptotic functions previously ascribed to *vif, env*, etc. (51, 52). As a cancer cell line, HUT78 cells are relatively resistant to apoptosis; this resistance is a result of null mutations in the *p53* gene (53, 54). However, since Vpr-induced apoptosis is p53-independent (25, 55), we believe that R77Q Vpr is a *bona fide* pro-apoptotic mutation while WT and R36W Vpr induce necrotic cell death. We saw an increased number of necrotic cells in the R36W-infected cells compared to the WT-infected population, however this could be a result of R36W’s enhanced replication capacity, and not due to enhanced cytotoxicity.

HIV has been shown to induce pyroptosis in bystander cells (56), but this process is mediated by cell-to-cell contact (57), primarily within three-dimensional lymphoid tissue models and not in peripheral blood cells (36, 58). Therefore, it is unlikely that the form of necrotic cell death observed in our suspension of T cells was pyroptosis (53). However, an apoptosis-independent form of HIV-1-mediated cell death has been previously reported (59, 60). More recently, necroptosis, a form of programmed necrotic cell death was shown to take place in HIV-1 infected cells (61). Due to similarities between our data and the data reported by Pan et al., we believe that the necrotic cell death observed in the WT, R36W, and Null populations was likely a result of necroptosis. The mechanism by which HIV-1 induces necroptosis is still unknown, but it is likely induced by a combination of host and viral factors and not directly induced by a specific viral protein (61). It is interesting to note however, that Pan et al. reported an inverse relationship between the levels of apoptosis and necroptosis in infected populations of cells, suggesting that the pathways serve as alternatives to one another. Mutations like R77Q that lead to a preference of one pathway over another could have a significant impact on host-virus interactions.

*In vivo*, a tendency to induce pro-inflammatory death could potentially lead to a faster and/or more potent chronic activation of the immune system, which has been hypothesized to lead to AIDS progression (62). Thus, the R36W mutation could show an RP phenotype based upon the avoidance of apoptosis and reliance on necrotic death mechanisms, which are highly inflammatory (63). This mechanism would fit in well with the RP phenotype reported by others (9). On the other hand, it is known that apoptosis does not enhance a systemic inflammation (64) and therefore could delay the chronic activation of the immune system that leads to AIDS. If the R77Q mutation influences host cells to die primarily by apoptosis, this could potentially explain the LTNP phenotype associated with it. Although R77Q is highly cytotoxic and replicates at a similar rate to the WT virus, the death pathway followed by infected cells could significantly alter the host’s immune response during chronic HIV-1 infection. Only an *in vivo* study, however, will be able to conclusively determine the progression phenotypes of the R36W and R77Q mutations in a more physiological setting.

In summary, we present novel replication and cytopathic phenotypes associated with the R36W and R77Q Vpr mutations that could potentially contribute to the overall role that HIV Vpr plays in CD4+ T cell depletion and disease progression.

## Materials and Methods

### Plasmids

HIV-1 pNL4-3 was obtained through the NIH AIDS Reagent Program, Division of AIDS, NIAID, NIH: HIV-1 NL4-3 Infectious Molecular Clone (pNL4-3) from Dr. Malcolm Martin (Cat# 114)(65–67). HIV-1 NL4-3 R36W Vpr and HIV-1 NL4-3 R77Q Vpr mutants were a gift from Dr. Velpandi Ayyavoo (University of Pittsburgh). The region surrounding the *vpr* gene was cloned into the pUC19 vector (Addgene #50005) using the EcoRI and SbfI sites (New England Biolabs) for SDM. SDM was accomplished using back to back primers with single point mismatches to change the ATG codon to GTG for the null mutant in order to not affect any overlapping genes in the viral genome (Forward: CAGAGGACAGGTGGAACAAGC; Reverse: TCAGTTTCCTAACACTAGGC). After SDM the *vpr* gene was removed from pUC19 using the same enzymes and ligated back into its original frame in the pNL4-3 vector. The desired mutation and in-frame ligations were confirmed by Sanger sequencing. One shot Stbl3 *E. coli* cells (Invitrogen C737303) were used for transformations to avoid plasmid recombination.

### Cell culture

HEK293FT cells were maintained at 37°C, 5% CO_2_ in DMEM (Sigma-Aldrich) supplemented with 10% fetal bovine serum (FBS; HyClone), 2% glutamine, 1% G418 (Teknova) and 1% penicillin/streptomycin (P/S). HUT78 cells were maintained at 37 °C, 5% CO2 in RPMI (Mediatech) supplemented with 10% FBS, 1% glutamine and 1% P/S. Ghost R3/X4/R5 cells were maintained at 37°C, 5% CO2 in DMEM, supplemented with 10% FBS, 1% P/S, 500 μg/mL G418, 100 μg/mL hygromycin, and 1 μg/mL puromycin. (Ghost cells were obtained through the NIH AIDS Reagent Program, Division of AIDS, NIAID, NIH: GHOST (3) CCR3+ CXCR4+ CCR5+ Cells from Dr. Vineet N. KewalRamani and Dr. Dan R. Littman (cat# 3943)(68)).

### Transfection and viral titration

HEK293FT cells were transfected using the calcium phosphate method; virus was collected 48 hours post-transfection. Viral concentration was determined by titration using Ghost R3/X4/R5 cell line using the methods described at the NIH AIDS Reagent Program (69).

### HIV-1 infections and flow cytometry analysis

Cells were infected with 0.1 MOI (cell cycle analysis) or 0.01 MOI (all other experiments) with polybrene. Cells were collected on the specified day of infection for each test and prepared as follows for Annexin V flow cytometry analysis. Cells were washed twice with serum-free PBS and centrifuged for 5 minutes at 500 x g, stained with Fixable Viability Dye (FVD) for 30 minutes at 4°C, and then washed with FACS staining buffer (FCSB) (140mM NaCl, 4mM KCl, 0.75 mM MgCl2, 10mM HEPES, 1% BSA, 0.1% NaN_3_), CaCl_2_ was then added at a concentration of 1.5mM for 10min before staining with fluorochrome-conjugated Annexin V for 15m at 25°C. Cells were then washed with FCSB and 1.5 mM CaCl_2_, fixed with Solution A and 1.5 mM CaCl_2_ for 30 minutes, washed with FCSB and permeabilized with Solution B (Fix & Perm, Nordic MUbio, GAS-002). Anti-p24 antibody was then added at a final dilution of 1:400, and cells were incubated for 15m at 25°C. Cells were then washed twice with FCSB, resuspended, and analyzed by flow cytometry using a Beckman Coulter Cytoflex Cytometer (70, 71). Results were analyzed using FlowJo software (version 10.6.2). The dyes used were: KC57 anti-HIV-1 core antigen-PE (Beckman coulter CO604667), Live/Dead fixable Far Red Dead Cell Stain Kit (Invitrogen L10120) and Annexin V FITC conjugate (Invitrogen A13199). For cell cycle analysis we used the FxCycle PI/ RNase Staining Solution (Invitrogen F10797) together with KC57 anti-HIV-1 core antigen-FITC (obtained through the NIH AIDS Reagent Program, Division of AIDS, NIAID, NIH: Anti-HIV-1 p24 Monoclonal (KC57)-FITC from NIAID, DAIDS (cat# 13450)).

For TUNEL assays, cells were stained with FVD, and then stained for TUNEL analysis using FragEL^™^ DNA Fragmentation Kit, Fluorescent-TdT Enzyme (Millipore QIA39) following the manufacturer’s protocol.

### Immunoblotting

HUT78 cells were washed with ice-cold PBS and lysed in RIPA buffer (ThermoFisher) with protease inhibitor cocktail (ThermoFisher). Proteins were quantified using BSA Protein Assay Kit (ThermoFisher). Equal amounts of protein were boiled with 6X loading buffer (375mM Tris HCL, 9% SDS, 50% Glycerol, 0.03% Bromophenol blue) for 10 minutes. Samples were run through a 4-15% gradient polyacrylamide gel and then transferred to a PVDF membrane (BioRad). The membrane was blocked with PBS-T containing 5% non-fat milk for 1h at 25°C. Vpr was probed using Anti-HIV-1 Vpr monoclonal (diluted 1:200) (Cosmo Bio) and Goat anti-Mouse HRP-conjugated (diluted 1:5,000) (Invitrogen) antibodies. Fibronectin was probed using Rabbit anti-Fibronectin (diluted 1:2,000) (Abcam) and Goat anti-Rabbit HRP-conjugated (1:20,000) (Abcam) antibodies. Detection was done using WesternBright ECL (Advansta) and results were analyzed using ImageJ software.

### Viral replication

Viral supernatants were collected on days 3, 5 and 7, and viral RNA was extracted using Viral Nucleic Acid Extraction kit II (Scientific FF10616-GB). The extracted RNA was used as template to produce cDNA using High Capacity cDNA Reverse Transcription Kit (Applied Biosystems 4368814) and LTR-specific primers. Finally, 2X Forget-Me-Not Universal Probe Master Mix with ROX (Biotium 31044-1) was used to perform Q-RT-PCR in an Applied Biosystems StepOnePlus^™^ Real-Time PCR System Thermal Cycling Block, following the protocol described previously by Rouet et al (72). The software used to analyze the data was StepOnePlus Software 2.2.3.Ink and results were normalized to RNA copies x mL^-1^.

### Statistical analysis

Biological samples were analyzed in triplicate using one-tailed Student’s t-test with a significance level of p ≤; 0.05.

## Acknowledgments

We would like to acknowledge Evelyn Wong for help with immunoblotting procedures, and Preston Neff and Brent Palmer for assistance with flow cytometry protocols.

